# Pangenome analysis provides insights into the evolution of mango (*Mangifera* spp.)

**DOI:** 10.1101/2025.05.09.653124

**Authors:** Jin Li, Vincent N. Michael, Sheron A. Simpson, Ramey C. Youngblood, Brian E. Scheffler, Jonathan Crane, Sisi Chen, Amanda M. Hulse-Kemp, Xingbo Wu

## Abstract

Mango (*Mangifera* spp.) is a major tropical fruit crop of global economic importance, but advanced genomic resources are needed to support its breeding, conservation, and sustainable cultivation. In this study, a mango pangenome was constructed using high-quality, telomere-to-telomere genomes of four cultivated mango (*M. indica*) accessions representing distinct genetic origins, and one wild relative, *M. odorata*. Genome-wide analyses revealed a significant reduction in heterozygosity among elite commercial cultivars, indicating a genetic bottleneck resulting from long-term artificial selection. Core genes were enriched in fundamental biological pathways, including primary metabolism, photosynthesis, transcriptional regulation, and cellular signaling. Variable genes were primarily associated with secondary metabolite biosynthesis, reflecting local environmental adaptations. Pangenome and comparative genomic analyses identified structural variations among accessions. Additionally, 10,482 high-confidence single nucleotide polymorphisms (SNPs) were detected and utilized for population genomic analysis of 197 mango accessions, delineating four genetically distinct groups. Southeast Asian accessions exhibited unique genetic diversity and divergence from Caribbean, Indian, and U.S. groups. Comparative analyses revealed differentiation in specialized metabolic pathways, particularly alkaloid and diterpenoid biosynthesis, likely reflecting adaptive responses to the complex ecological interactions and high biodiversity of Southeast Asian tropical rainforests. Genomic analysis of the *MiRWP* gene, associated with apomixis, provided comprehensive insights into this important mango trait, demonstrating the potential of the pangenome for future mango breeding efforts. The genomic resources generated in this study establish a critical foundation for advancing mango genetic research, facilitating trait improvement, and informing conservation strategies.

## Introduction

Mango (*Mangifera indica* L.) is a major tropical fruit crop renowned for its distinctive aroma, sweet flavor, and rich nutritional properties, including polyphenolics and vitamins A, C, and E (Lauricella et al., 2017; Liu et al., 2020). Often referred to as the “king of fruits”, mango has experienced rising global demand, with total production exceeding 59.15 million tons in 2022 (Shahbandeh, 2024). Major producers include India, Indonesia, China, Pakistan, Mexico, and Brazil (World Population Review, 2024). In the United States, mango consumption has increased significantly over the past two decades, with per capita intake rising from 1.75 pounds in 2000 to 3.66 pounds in 2021 (Shahbandeh, 2023).

The genus *Mangifera* (family Anacardiaceae) comprises over 60 species (Bompard, 2009), with *M. indica* being the primary species cultivated commercially. Other species, including *M. odorata* Griff., *M. foetida* Lour., *M. pajang* Kosterm., and *M. caesia* Jack, also produce edible fruits (Kostermans and Bompard, 1993; Prasad et al., 2022). Extensive *Mangifera* germplasm collections, particularly of *M. indica*, are maintained across subtropical and tropical regions worldwide (Bettencourt et al., 1992). Key global repositories, including those in India, Australia, China, Israel, and the United States, play a crucial role in conserving *Mangifera* genetic diversity and supporting breeding efforts (Dillon et al., 2013; Sherman et al., 2015; Souza et al., 2020; Liang et al., 2024; Ali et al., 2025; Eltaher et al., 2025). In the United States, there are two major mango collections: one maintained by the United States Department of Agriculture (USDA) Plant Germplasm Bank and the other preserved at the University of Florida Tropical Research and Education Center (UF-TREC). While the USDA-ARS collection represents a wide genetic diversity of mango species, the UF-TREC mango collection comprises popular cultivars that resulted from human selection of chance seedlings.

Mango seedlings typically require a minimum of five years to produce their first fruit. This prolonged juvenile phase poses a major challenge for trait evaluation and significantly constrains mango breeding. Elucidating the genetic diversity and evolutionary history of *Mangifera* species provides essential insights into adaptive traits and fruit quality, thereby facilitating breeding programs, market expansion, and biodiversity conservation.

Fossil evidence indicates that *Mangifera* originated in Peninsular India during the Paleocene, with Southeast Asia emerging as its principal diversification center, which is supported by the high species richness of the region (Mehrotra et al., 1998). Historically, it was postulated that mango underwent a single domestication event in India over 4,000 years ago, subsequently spreading to Southeast Asia during the 4th and 5th centuries CE (Mukherjee, 1950; Mehta, 2017; Warschefsky and von Wettberg, 2019), and then dispersed globally from the 14th century onward (Mukherjee, 1972; Mehta, 2017). A more complex domestication history has also been suggested in which there were independent domestication events in both India and Southeast Asia. This hypothesis is supported by the higher genetic diversity observed in Southeast Asian mangoes, alongside distinct differences in fruit morphology and embryony type when compared to Indian mangoes (Iyer and Schnell, 2009; Warschefsky and von Wettberg, 2019; Wang et al., 2020; Fairchild Tropical Botanic Garden, n.d.). Recent population genomic studies utilizing high-quality SNP markers further substantiate this hypothesis, delineating Indian and Southeast Asian mangoes into two distinct genetic clusters (Warschefsky and von Wettberg, 2019; Wang et al., 2020; Liang et al., 2024; Ma et al., 2024). U.S. cultivars have been shown to exhibit a closer genetic relationship to Indian mangoes, suggesting a shared ancestral lineage (Warschefsky and von Wettberg, 2019; Wang et al., 2020; Ma et al., 2024).

Mango possesses a diploid genome (2n = 2x = 40) with a compact genome size ranging from approximately 365 Mb to 439 Mb, depending on the cultivar (Li et al., 2020; Wang et al., 2020; Bally et al., 2021; Wijesundara et al., 2024). Mango’s high heterozygosity, resulting from outcrossing and hybridization, presents challenges for high-quality genome assembly (Mukherjee, 1950). Several mango genomes of varying quality are available, capturing genetic variation at the single-accession level (Singh et al., 2016; Wang et al., 2020). Advances in long-read sequencing technologies, such as PacBio HiFi and Revio (>99.9% accuracy, 10-25 kb reads), have improved the feasibility of assembling complex genomes (Hon et al., 2020; Li and Cullis, 2023; Li and Cullis, 2025). The first telomere-to-telomere (T2T) high-quality genome assembly of *M. indica* (cultivar ‘Irwin’) was generated using PacBio HiFi sequencing, with contigs anchored to the *M. indica* ‘Alphonso’ reference genome, followed by manual curation (Wang et al., 2020; Wijesundara et al., 2024).

In contrast to single-genome assemblies, pangenome analyses offer a comprehensive perspective of genetic diversity across accessions, encompassing both core genes shared by all individuals and accessory genes that exhibit variability among varieties. These analyses also provide insights into how structural variations (SVs) can influence gene expression. Structural variations (SVs), such as insertions and deletions, can disrupt regulatory regions or coding sequences, directly altering gene activity. Gene duplications may result in increased gene dosage, potentially enhancing expression levels, while gene fragmentations can compromise gene integrity, leading to reduced or silenced expression (Alonge et al., 2020). Collectively, these variations contribute to genetic diversity and influence heritable traits (Tay Fernandez et al., 2022). Pangenome analyses have been applied to various fruit crops, including tomato (*Solanum lycopersicum*) (Gao et al., 2019; Zhou et al., 2022), apple (*Malus domestica*) (Wang et al., 2023), orange (*citrus* spp.) (Huang et al., 2023), and strawberry (*Fragaria* spp.) (Qiao et al., 2021). Critical functional genes associated with key traits have been pinpointed in these studies, offering valuable breeding resources. For instance, genes related to fruit color have been identified in strawberry (Qiao et al., 2021), apple (Wang et al., 2023), and watermelon (*Citrullus lanatus*) (Zhang et al., 2024), while genes influencing flavor have been discovered in tomato (Gao et al., 2019; Zhou et al., 2022) and orange (Huang et al., 2023). Additionally, the pangenome has enabled the identification of genes conferring disease resistance in watermelon (Zhang et al., 2024).

In this study, T2T genome assemblies were generated for four mango cultivars: ‘Haden’, ‘Carabao’, ‘Alampur Baneshan’ (Alampur), and ‘Safeda Lucknow’ (Safeda), representing U.S., Southeast Asian, and Indian genetic diversity as demonstrated by Michael et al. (2023). Additionally, a mango accession of *M. odorata* was assembled to represent the wild species. Pangenome analyses were performed to identify core and variable gene families across these accessions, followed by enrichment analyses to highlight pertinent gene pathways. Major genomic structural variations were assessed through whole-genome sequence alignment. Evolutionary relationships were explored by constructing a phylogenetic tree based on shared orthologs and analyzing potential whole-genome duplication (WGD) events. Furthermore, a total of 10,482 high-quality SNPs were utilized to assess the genetic diversity and population structure of 197 *Mangifera* accessions maintained at the UF-TREC, followed by an investigation of genome-wide selection within sub-populations. The genomic resources generated in this study provide valuable insights into the evolutionary dynamics of the *Mangifera* genus and offer an important foundation to accelerate mango breeding efforts.

## Results

### Genome assembly and annotation of mango accessions reveal high-quality genomic features and evolutionary insights

High-quality PacBio HiFi sequencing was performed for all selected accessions, with sequencing outputs ranging from 21.0 Gb to 34.3 Gb (on average 50x genome coverage) (Supplementary Table S1). Genome assemblies were generated using HiFiasm v.0.19.9 (Cheng et al., 2021) and scaffolded with RagTag v.2.0.1 (Alonge et al., 2022), using the T2T *M. indica* ‘Irwin’ genome as reference (Wijesundara et al., 2024). All five assemblies reached T2T chromosome-scale resolution, with genome sizes spanning from 365.73 Mb (‘Alampur’) to 368.46 Mb (‘Carabao’) and scaffold N50 values ranging from 18.43 Mb (‘Carabao’) to 19.17 Mb (*M. odorata*) (Table 1).

**Table 1.**
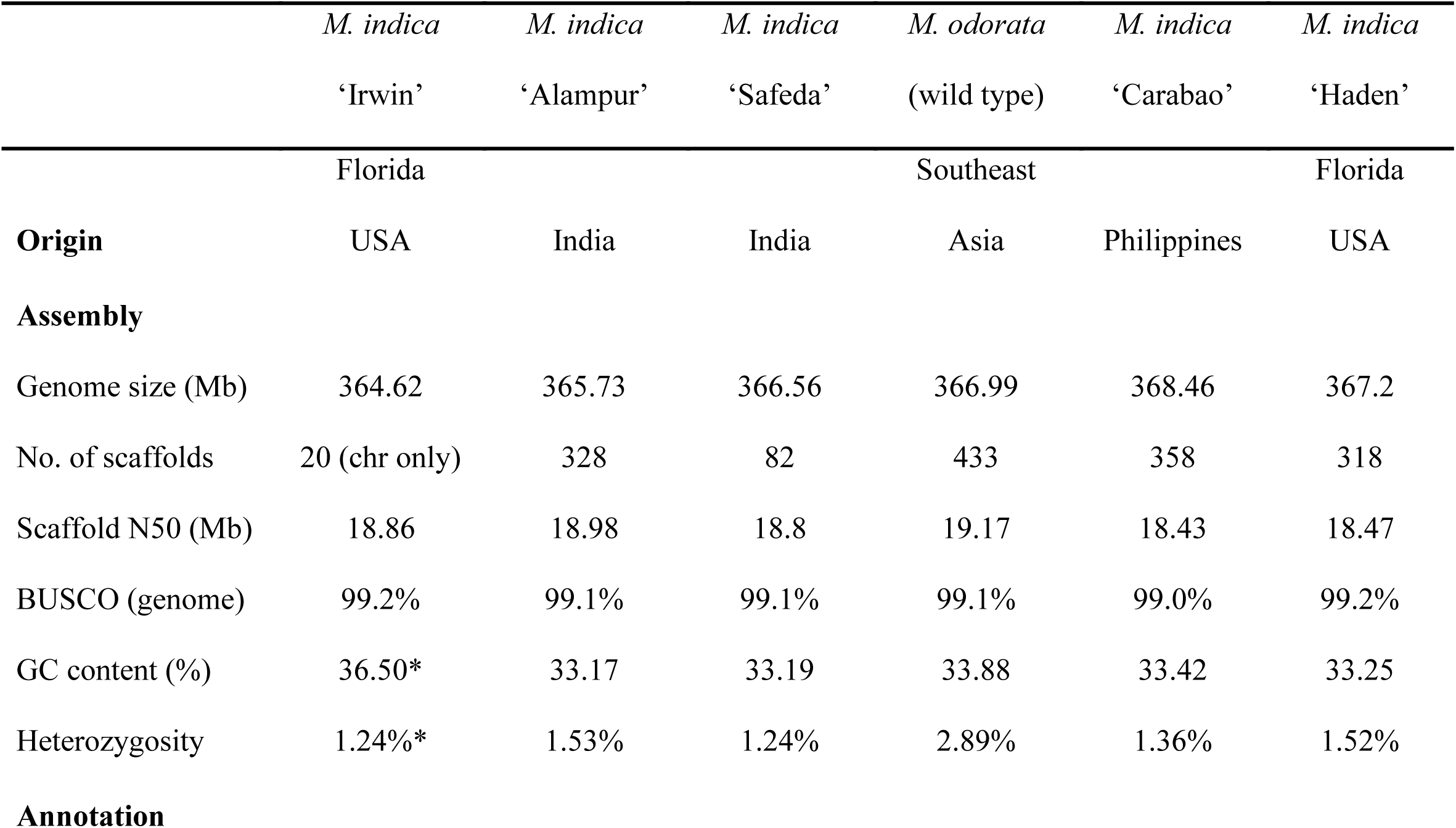
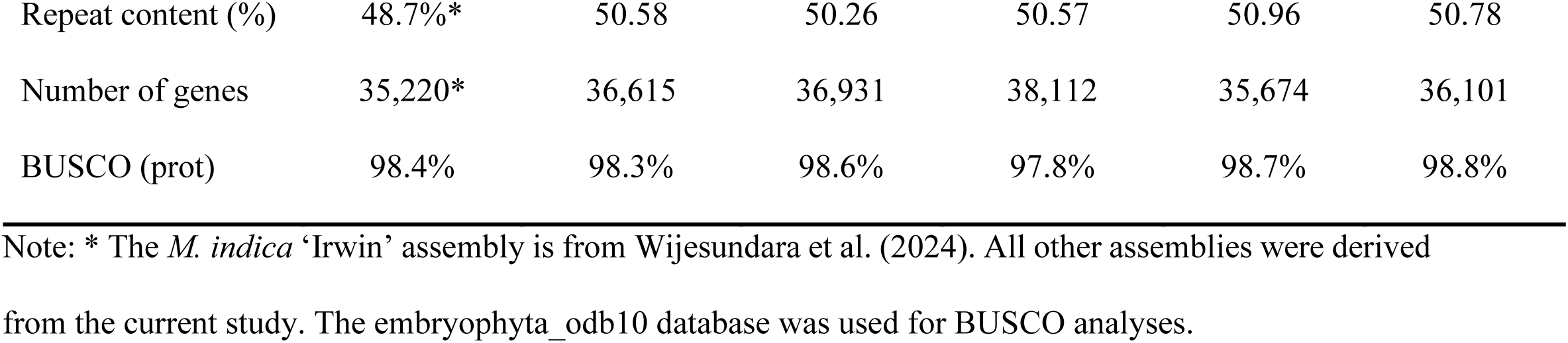
Genomic features and quality metrics of six Mangifera primary haplotype assemblies.

The genome assembly completeness of all five accessions exceeded 99% based on the Benchmarking Universal Single-Copy Orthologs (BUSCO) v.5.3.0 (Manni et al., 2021), using the embryophyta_odb10 database. The GC content of the five accessions ranged from 33.17% to 33.88%, lower than that of the collapsed ‘Irwin’ genome (36.50%) (Wijesundara et al., 2024). The highest heterozygosity (2.89%) was observed in *M. odorata*; the domesticated accessions ranged from 1.24% (‘Safeda’ and ‘Irwin’) to 1.53% (‘Alampur’). Repetitive elements accounted for 50.26% to 50.96%, with long terminal repeats (LTRs) being the most abundant, contributing 17.50% to 18.51% of the assembled genomes (Supplementary Table S2).

Gene prediction was performed using the BRAKER v.3.0.8 pipeline “C” (Gabriel et al., 2024), integrating protein-spliced alignments and start/stop codons signals. Orthologous groups from OrthoDB v.12.0 (Kuznetsov et al., 2022) and reference proteins from ‘Irwin’ were used as inputs for training. The total predicted genes ranged from 35,674 (‘Carabao’) to 38,112 (*M. odorata*), with a minimum BUSCO score of 97.8% (Table 1). Structural annotation revealed that the *M. odorata* genome has an average of 5.79 exons (mean size: 222.68 bp) and 4.67 introns (mean size: 369.67 bp) per gene, whereas ‘Carabao’ has an average of 6.22 exons (mean size: 219.10 bp) and 5.09 introns (mean size: 368.72 bp).

### Pan-genome characterization and functional insights across six mango accessions

OrthoFinder identified 33,839 orthogroups with 224,357 genes across the six *Mangifera* accessions (Figure 1). Of these, 22,592 orthogroups (66.8%) with a total of 172,777 genes (77.0%) were classified as core gene families shared across all accessions (Figure 1a). An additional 3,435 orthogroups (10.2%), comprising 22,878 genes (10.2%), were designated as softcore gene families shared by five accessions. A total of 626 orthogroups (1.8%), containing 1,929 genes (0.9%), were exclusive to *M*. *odorata*. The number of accession-specific orthogroups ranged from 42 in ‘Haden’ to 260 in ‘Irwin’. The total number of gene clusters increased until reaching a plateau, indicating the constructed pangenome captures the majority of the gene content within *Mangifera* accessions. Concurrently, the proportion of core clusters declined to approximately two-thirds of the total clusters (Figure 1b). Core orthogroups constituted the highest proportion of total orthogroups in ‘Irwin’ (80.99%) and the lowest in ‘Safeda’ (75.90%), whereas exclusive clusters accounted for 0.14% of total clusters in ‘Haden’ and 0.93% in ‘Irwin’ (Figure 1c, Supplementary Table S3).

**Figure 1.**
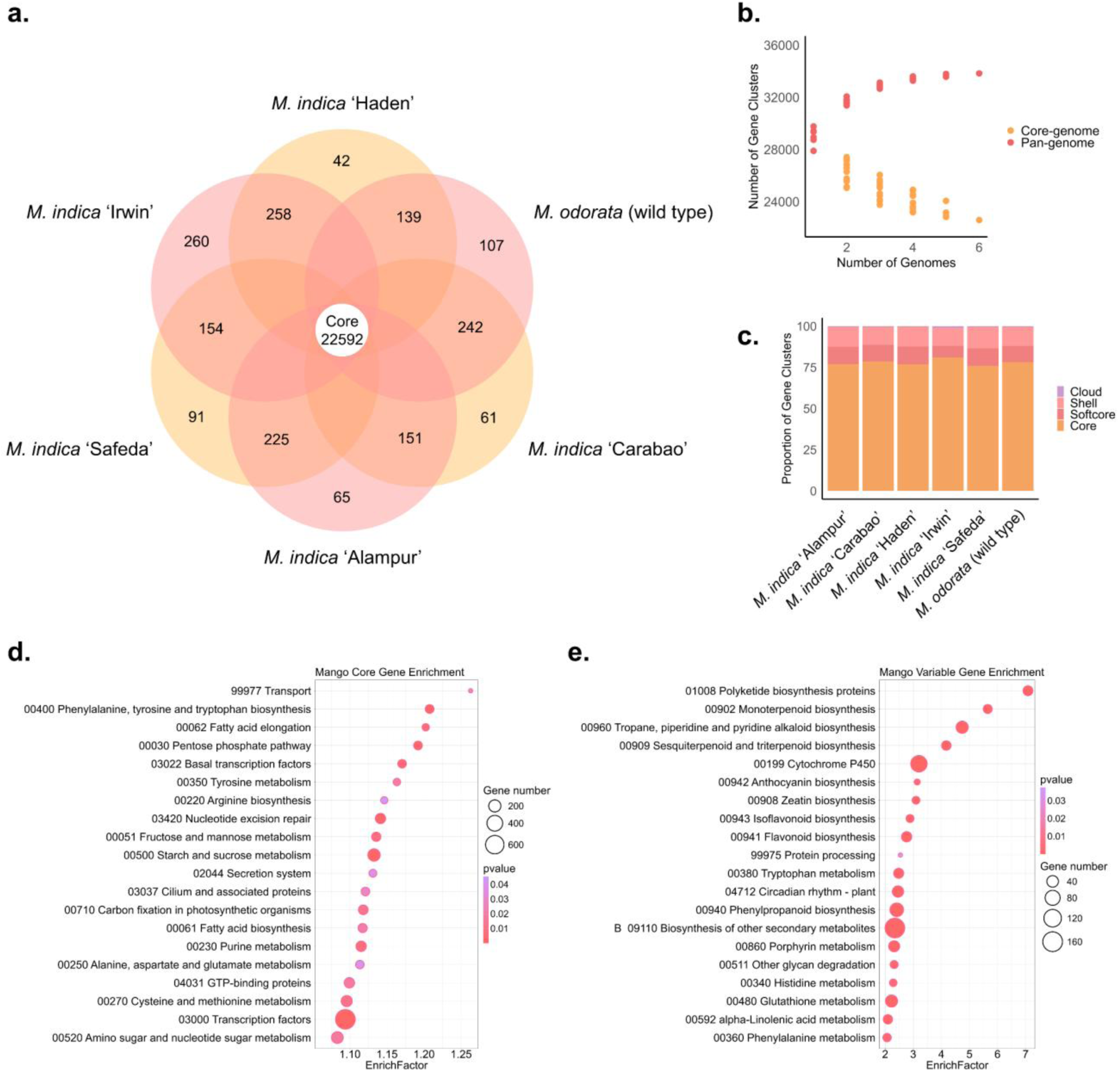
Pan- and core-genome analyses of six mango accessions. (a) Venn diagram illustrating the number of core, accession-specific, and shared gene families among the six *Mangifera* accessions. (b) Variation in gene family numbers across the pan-genome and core-genome, along with additional *Mangifera* genome assemblies. (c) Stacked histogram showing the proportion of core, soft core (shared by five out of six accessions), shell (moderately conserved genes in two to four accessions), and cloud (accession-specific) gene families in *Mangifera* accessions. (d) KEGG pathway enrichment analysis of genes in core gene families. (e) KEGG pathway enrichment analysis of genes in variable gene families (soft core, shell, and cloud).

Kyoto Encyclopedia of Genes and Genomes (KEGG) enrichment analysis revealed that core genes were significantly enriched in pathways associated with primary metabolism, genetic information processing, cellular signaling, and photosynthesis (Figure 1d, Supplementary Table S4). Notable enriched pathways include carbohydrate metabolism (e.g., starch and sucrose metabolism, pentose phosphate pathway), lipid biosynthesis and elongation, amino acid biosynthesis and metabolism, and nucleotide metabolism. Enrichment in photosynthesis-related pathways, such as carbon fixation, further emphasizes the importance of primary productivity in mango (Stirbet et al., 2019). Collectively, these pathways establish the molecular foundation for mango growth, development, and adaptation.

In contrast, variable genes were predominantly enriched in secondary metabolism pathways, such as the biosynthesis of terpenoids, flavonoids, alkaloids, and phenolic compounds (Figure 1e, Supplementary Table S5). These pathways are essential for the production of specialized metabolites involved in plant defense, pigmentation, and ecological interactions (Zaynab et al., 2018; Divekar et al., 2022; Elshafie et al., 2023; Tang et al., 2024). Additional enrichments were observed in protein processing, cytochrome P450 activity, and glutathione metabolism, contributing to enzymatic functions, detoxification, and antioxidant defenses (Xu et al., 2015; Dorion et al., 2021). Pathways related to circadian rhythm regulation suggest adaptations to environmental fluctuations (Venkat and Muneer, 2022; Xu et al., 2023). These findings highlight the metabolic and functional diversity within *Mangifera* accessions, which can drive phenotypic variation and environmental resilience.

### Comparative genomic analysis and evolutionary relationships of Mangifera accessions reveal structural variation, divergence, and adaptation

The phylogeny of Sapindales species, inferred from 1,820 shared single-copy orthologs, provides insights into their evolutionary relationships. The estimated divergence between *Mangifera* and *Pistacia* occurred approximately 34.36 million years ago (Ma), while *Mangifera* and *Citrus* diverged around 78 Ma (Figure 2a). Within *Mangifera*, phylogenetic analysis based on 14,243 concatenated single-copy orthologs revealed a similar topology, with Indian cultivars (‘Alampur’ and ‘Safeda’) forming early-diverging clades. Notably, ‘Alampur’ clusters closely with U.S. cultivars (‘Irwin’ and ‘Haden’), despite substantial genetic differentiation, suggesting localized adaptation and prolonged genetic isolation. Gene family variation analysis indicated that gene family contractions generally outnumber expansions, suggesting genome simplification in most accessions. However, ‘Alampur’ exhibited more expansions than contractions, potentially due to adaptation to diverse environments or retention of ancestral genomic features.

**Figure 2.**
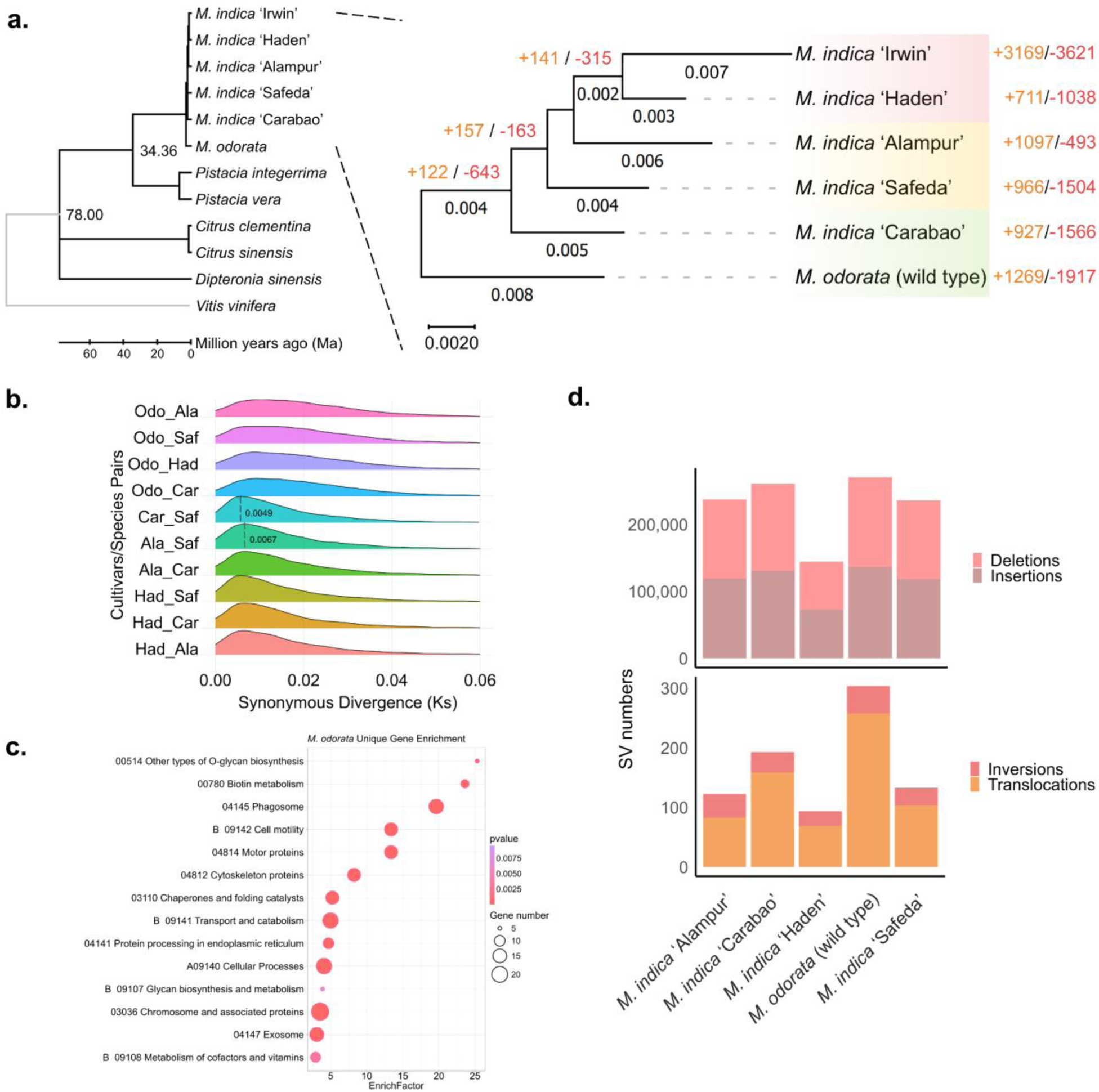
Comparative genomic analysis of *Mangifera* accessions. (a) Maximum likelihood phylogenetic trees of six *Mangifera* accessions and five other species from the order Sapindales (*Pistacia vera*, *Pistacia integerrima*, *Citrus clementina*, *Citrus sinensis*, and *Dipteronia sinensis*), with *Vitis vinifera* as the outgroup. Phylogenies were inferred from MAFFT alignments of concatenated shared single-copy orthologs identified by OrthoFinder. In the species-level tree (left), estimated divergence times (in millions of years) are labeled on the branches. In the *Mangifera* tree (right), the counts of expanded and contracted gene family are indicated next to the accession names and branches. Shading beneath accession names denotes geographical origins: red for Florida-USA, yellow for India, and green for Southeast Asia. (b) Pairwise synonymous substitution rate (Ks) distribution derived from alignments of shared single-copy orthologs. Accession names are abbreviated using the first three letters for simplicity. (c) KEGG pathway enrichment analysis of genes in *M. odorata-*exclusive gene families, ranked by enrichment factor, with dot size representing the number of genes. (d) Structural variation (SV) analysis of five *Mangifera* genomes aligned to the ‘Irwin’ reference genome.

The distribution of synonymous substitutions per site (Ks) was calculated for orthologous gene pairs to further estimate divergence among the six mango accessions (Figure 2b). *M. odorata* showed a broader Ks distribution and earlier divergence from domesticated *M. indica* cultivars, reflecting long-term evolutionary separation. This extended Ks profile highlights *M. odorata* as a potential reservoir of genetic diversity for breeding, particularly for traits such as disease resistance and environmental resilience. Among domesticated cultivars, ‘Alampur’ and ‘Safeda’ exhibited the earliest divergence (Ks = 0.0067), despite both originating from India. In contrast, ‘Carabao’ (Southeast Asian type) and ‘Safeda’ (Indian type) showed a more recent divergence (Ks = 0.0049). KEGG enrichment analysis of 7,828 genes from 107 *M. odorata*-exclusive gene families (Figure 2c, Supplementary Table S6) revealed significant enrichment in pathways related to cellular structure, protein processing, and specialized metabolism. Enriched pathways associated with the cytoskeleton, motor proteins, and cell motility suggest enhanced cellular organization and transport mechanisms (Wang and Mao, 2019; Motta and Schnittger, 2021; Schramma et al., 2022). Enrichment of protein processing pathways, particularly those related to the endoplasmic reticulum and chaperone activity, indicates improved proteostasis supporting stress tolerance and cellular functionality (Gupta and Tuteja, 2011; Reyes-Impellizzeri and Moreno, 2021). Additionally, enrichment in glycan biosynthesis and biotin metabolism pathways highlights specialized metabolic processes contributing to environmental adaptability (Varki, 2016; Chaliha et al., 2018; Wang et al., 2020).

Genome synteny and structural variation analyses revealed substantial genomic divergence among the *Mangifera* accessions (Figure 3, Supplementary Table S7). Syntenic regions accounted for 91.49% (*M. odorata*, 335.77 Mb) to 97.09% (‘Haden’, 356.50 Mb) of the assemblies, reflecting their phylogenetic proximity to the reference genome ‘Irwin’. Structural variations, including insertions, deletions, inversions, translocations, and duplications, were detected, with insertions and deletions being the most abundant types. The number of insertions ranged from 72,671 to 136,366, covering 1.23 Mb to 1.69 Mb, while deletions ranged from 71,794 to 134,240, covering 1.25 Mb to 1.72 Mb across the genomes. Among the genomes, ‘Haden’ exhibited the fewest, and *M. odorata* the most structural variation. Inversions were notably higher in *M. odorata* (46 inversions spanning 4.19 Mb) compared to ‘Haden’ (25 inversions spanning 2.51 Mb). *M. odorata* also showed the highest number of translocations (258 events spanning 3.29 Mb) and duplications (328 events spanning 1.95 Mb), indicating extensive genome restructuring and expansion. Unaligned genomic regions, reflecting structural divergence or assembly gaps, were most pronounced in *M. odorata*, consistent with its high sequence divergence. A similar trend was observed at the nucleotide level, where *M. odorata* displayed the highest SNP density (3.00 million SNPs), while ‘Haden’ exhibited the lowest (1.52 million SNPs). These findings underscore the genomic complexity and evolutionary differentiation among *Mangifera* accessions, particularly highlighting *M. odorata* as a wild species harboring pronounced structural and sequence-level variations potentially underlying its distinct adaptive and phenotypic traits.

**Figure 3.**
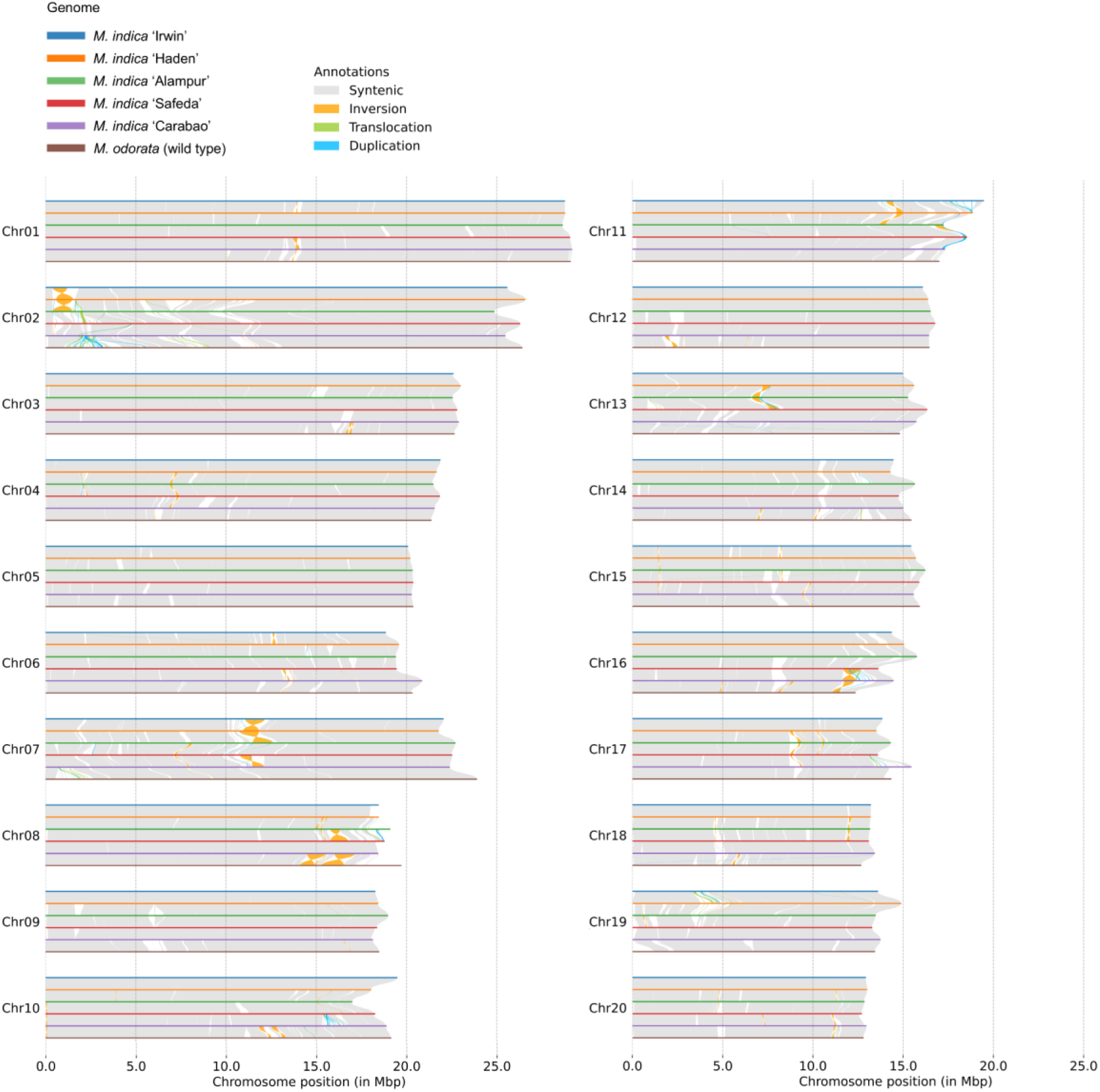
Genome-wide pairwise synteny and structural variation analysis of five *Mangifera* assemblies compared to the reference genome ‘Irwin’. The synteny plot shows conserved genomic regions (in grey) and structural variations (inversions, translocations, and duplications) across 20 chromosomes, identified using SyRI. Structural variants are color-coded and annotated, highlighting genome rearrangements among accessions.

Pairwise alignments of *Mangifera* assemblies further revealed the distribution of major structural variations across the genome (Figure 3). Notably, a genomic region of chromosome 2 exhibited extensive structural differences, particularly between *M. odorata* and ‘Carabao’, despite both having a Southeast Asian origin. This region harbored widespread tandem duplications and divergent interspersed sequences. Tandem duplications were also observed on chromosomes 10, 11, and 16. Inversions were prevalent across multiple genomic regions. ‘Haden’ displayed a large inversion at the 5’ end of chromosome 2, which was absent in all other accessions. A unique inversion near the centromere of chromosome 7 distinguished ‘Carabao’ from ‘Safeda’. Two adjacent inversions were identified on chromosome 8 between *M. odorata* and ‘Carabao’, alongside a major inversion between ‘Alampur’ and ‘Safeda’. Additional large inversions on chromosomes 11, 13, and 16 further underscore the dynamic structural variation landscape within *Mangifera* genomes.

### Genome-wide nucleotide diversity and genetic structure of mango accessions

To investigate genome-wide nucleotide diversity, 10,482 high-quality SNPs were obtained via genotyping-by-sequencing (GBS) for 197 mango accessions maintained at UF-TREC (Supplementary Tables S8 and S9). ADMIXTURE analysis identified the optimal number of genetic clusters as *K* = 4 (Figure 4a, Supplementary Figure S1), corresponding to major geographic regions where mango is extensively cultivated: Florida-USA, India, the Caribbean, and Southeast Asia (Figure 4b).

**Figure 4.**
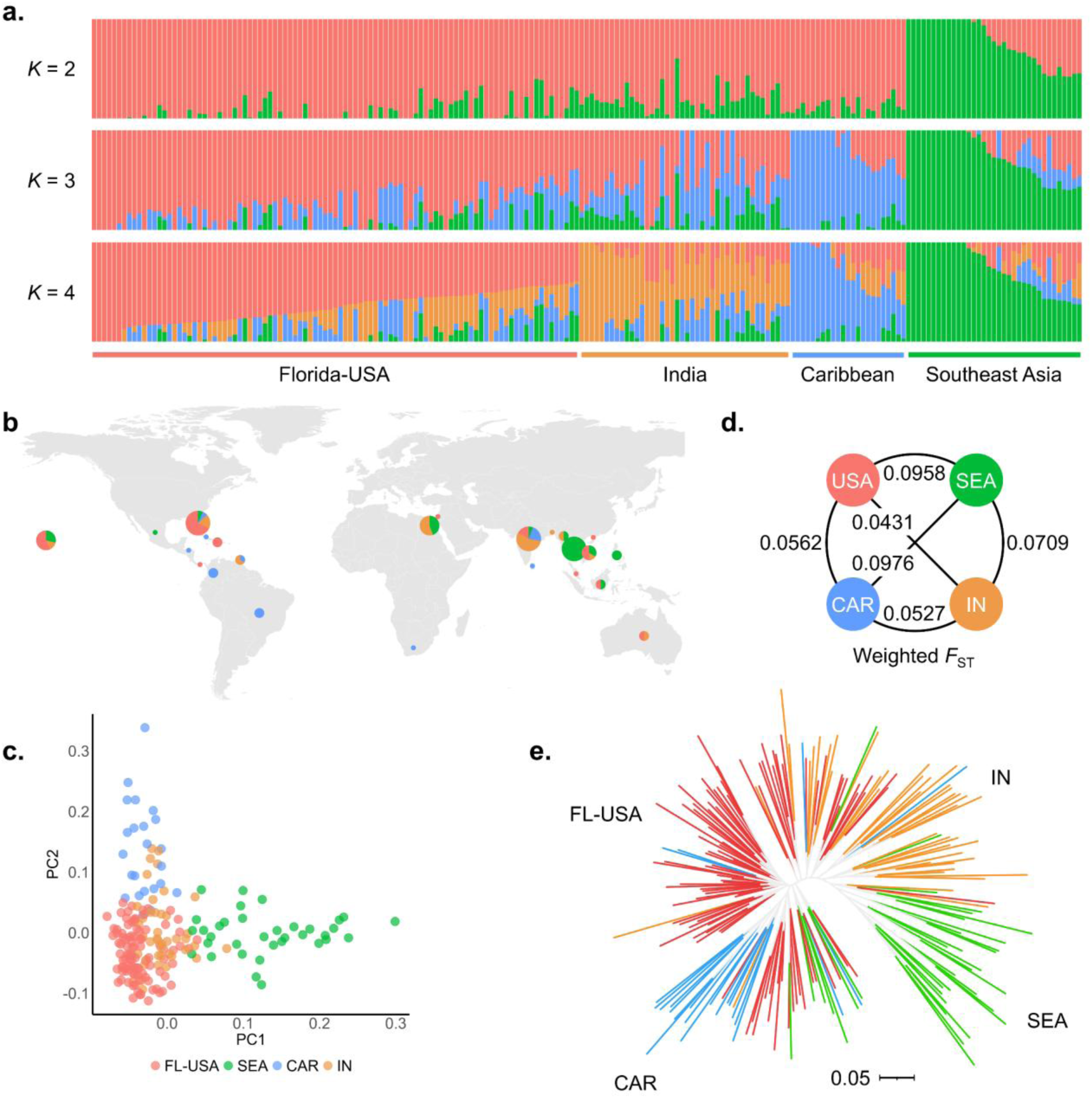
Comprehensive genomic population analysis of 197 *Mangifera* accessions. (a) Admixture analysis showing population structure and assignment probabilities based on 10,482 SNPs at *K* = 2, 3, and 4. At *K* = 4, the optimal number of clusters, determined by cross-validation, accessions are grouped into four distinct groups: Florida-USA (red), India (orange), Caribbean (blue), and Southeast Asia (green). (b) Geographical origin of accessions represented by pie charts, with chart size indicating the number of accessions per country. The proportions of the four population groups—Florida-USA (red), India (orange), Caribbean (blue), and Southeast Asia (green)—are depicted within each nation, illustrating the global distribution and genetic composition. (c) Principal component analysis (PCA) plot based on the top two principal components, highlighting the genetic structure and diversity among the 197 *Mangifera* accessions, with clustering patterns reflecting population differentiation. (d) Pairwise weighted F-statistics (*F_ST_*) analysis among the four population groups, quantifying genetic differentiation and population divergence. (e) Unrooted maximum likelihood (ML) tree depicting phylogenetic relationships among the accessions.

Genetic clustering revealed dynamic patterns of population structure across different *K* values. At *K* = 2, accessions from Florida, India, and the Caribbean grouped together. At *K* = 3, Caribbean accessions formed a separate cluster, indicating a distinct genetic background relative to Florida and Indian accessions. At *K* = 4, the Florida and Indian groups further diverged into individual clusters. In contrast, Southeast Asian accessions consistently formed a distinct group across all *K* values, suggesting a high degree of genetic differentiation.

Principal component analysis (PCA) provided further resolution of genetic relationships among accessions. Distinct clustering was observed for Florida-USA, Caribbean, and Southeast Asian groups, while the Indian group occupied an intermediate position, overlapping with both the Florida-USA and Caribbean groups (Figure 4c). This pattern suggests substantial genetic contributions from Indian mangoes to those regions. Pairwise *F_ST_* analysis indicated generally low genetic differentiation among the groups, with the highest *F_ST_* values remaining below 0.1 (Figure 4d). The Indian, Florida-USA, and Caribbean groups exhibited pairwise *F_ST_* values ranging from 0.043 to 0.056, reflecting minimal divergence. In contrast, the Southeast Asian group displayed the highest differentiation (*F_ST_* = 0.071 to 0.098) to the other three groups, reinforcing its distinct genetic lineage.

Phylogenetic analysis using an unrooted maximum likelihood (ML) tree further supported these findings, delineating the four genetic clusters while identifying several admixed accessions, likely resulting from hybridization or historical gene flow. Together, these results provide new insights into the genetic structure, evolutionary differentiation, and population history of cultivated and wild *Mangifera* accessions. The pronounced divergence of the Southeast Asian group highlights its unique genetic heritage, while the genetic connectivity among Indian, Florida-USA, and Caribbean groups reflects historical germplasm exchange and domestication processes.

Linkage disequilibrium (LD) decay patterns differed among the four mango subpopulations. Although all groups exhibited a general decline in *r*² values with increasing SNP distance (Supplementary Figure S2), the extent of LD decay varied. The Caribbean group had the highest average *r*² (0.0613), followed by Southeast Asia (0.0460), India (0.0359), and Florida-USA (0.0229). The Florida (USA) group exhibited the most rapid LD decay, likely reflecting its greater genetic diversity and larger sample size.

### Genomic selection signatures and evolutionary divergence between Indian and Southeast Asian mango groups

Due to low genetic divergence and high genomic similarity among mango accessions from the United States, India, and the Caribbean, and their shared historical origins linked to India, these three regional groups were merged into a single Indian group (IND) for comparative analyses with the Southeast Asian group (SEA). Genomic regions under selection were identified using SweeD, nucleotide diversity (log2(π _India_/ π _Southeast Asia_)), and *F_ST_* analyses (Figure 5, Supplementary Tables S10-S12), with a stringent 99.9th percentile threshold applied to detect strong selective signals and to minimize false positives.

**Figure 5.**
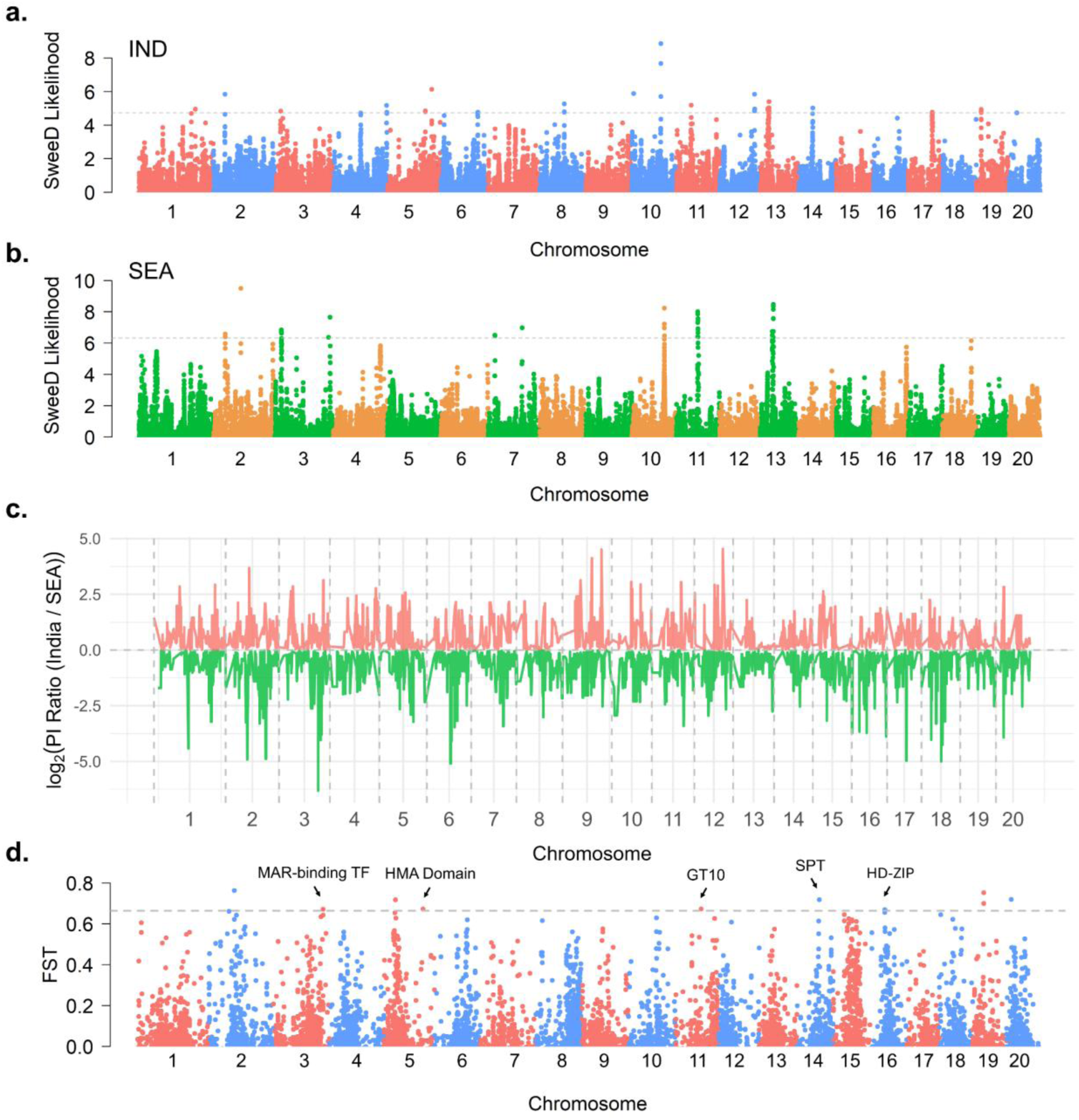
Genomic signatures of selection between combined Indian (Florida-USA, India, and Caribbean) and Southeast Asian *Mangifera* groups. (a) SweeD analysis of the combined Indian group (IND) reveals genome-wide signals of selective sweeps. (b) SweeD analysis of the Southeast Asian group (SEA) highlights regions under selection. SweeD likelihood values were calculated using SNPs in 10 kb windows, with the dashed line marking the 0.1% significance threshold. (c) Nucleotide diversity (π) was compared between the two groups using 200 kb sliding windows with 50-kb step sizes. The π ratio of the combined Indian group to the Southeast Asian group was log₂-transformed to emphasize genomic regions where diversity significantly deviates between the two groups, indicating potential ongoing selection pressures. (d) *F*_ST_ analysis identifies genomic regions of high differentiation between IND and SEA. The dashed grey line represents the 0.1% threshold (*F*_ST_ = 0.6640). Genes within the top *F*_ST_ regions include *MAR-binding TF* (transcription factor binding AT-rich DNA), *HMA Domain* (heavy metal binding), *GT10* (fucosyltransferase for cell wall biosynthesis), *SPT* (sugar phosphate translocator), and *HD-ZIP* (Homeodomain-leucine zipper).

SweeD analysis revealed distinct selective sweep patterns between the two groups. In the Indian group, significant selective sweeps were detected on chromosomes 5, 8, 11, 12, 14, 17, 19, and 20, whereas these regions did not show pronounced sweeps in the Southeast Asia group. Conversely, Southeast Asia-specific sweeps were observed on chromosomes 2 (mid-region), 3 (3’ end), 7, and 11.

Nucleotide diversity analysis further highlighted group-specific signatures of selection. The Indian group exhibited elevated diversity in regions of chromosomes 9 and 12, while the Southeast Asia group showed higher diversity on chromosomes 2, 3, 6, 17, and 18. These contrasting diversity patterns suggest localized selection and adaptation within each group.

The *F_ST_* analysis revealed strong genetic differentiation between Indian and Southeast Asian accessions across chromosomes 2, 3, 5, 11, 14, 16, 19, and 20. Many of these regions overlapped with SweeD-identified sweeps, underscoring their evolutionary significance. Genes located within these differentiated regions encode membrane proteins (e.g., At4g09580-like) and transporters, such as sugar phosphate translocators, which play roles in cellular transport and metabolic regulation (Dyson et al., 2014; Weise et al., 2019). The presence of stress-related proteins with heavy-metal-associated domains suggests adaptive responses to environmental challenges (Li et al., 2020). Regulatory divergence was also evident, as indicated by transcription factors including MAR-binding proteins and homeobox-associated leucine zippers (*HD-ZIP*), which are involved in gene expression regulation and developmental processes (Elhiti and Stasolla, 2009; Wang et al., 2010). Additionally, genes encoding chloroplastic proteins and glycosyltransferases point to differences in photosynthesis activity and secondary metabolism (Tiwari et al., 2016).

Notably, the clustered signal on chromosome 16, where the *HD-ZIP* gene resides, overlaps with a major GWAS signal for seed thickness in mango (Eltaher et al., 2025). Given the role of *HD-ZIP* transcription factors in regulating fruit size, seed number, and stress responses (Li et al., 2019; Li et al., 2022), this locus warrants further functional characterization.

KEGG enrichment analysis of genes within the top 5% *F_ST_* regions (200 kb windows) revealed pathways associated with specialized metabolism, transport, and cellular processes (Supplementary Figure S3, Supplementary Table S13). Key metabolic pathways included isoquinoline alkaloid biosynthesis, diterpenoid biosynthesis, and tropane, piperidine, and pyridine alkaloid biosynthesis, all of which are linked to bioactive compounds important for plant defense and environmental adaptation (O’Hagan, 2000, Matsuura and Fett-Neto, 2015; Li et al., 2021; Plazas et al., 2022). Variability in tyrosine and phenylalanine metabolism highlights differences in amino acid use and secondary metabolite synthesis (Tzin and Galili, 2010; Parthasarathy et al., 2018). Divergence in nicotinate and nicotinamide metabolism implies variation in energy and coenzyme biosynthesis (Hong et al., 2020), while sphingolipid metabolism may reflect differences in membrane composition and signaling (Luttgeharm et al., 2016). Transport-related pathways, such as ABC transporters and other membrane-associated systems, suggest group-specific nutrient uptake and molecular exchange mechanisms (Kang et al., 2011), while enrichment of glycan degradation pathways highlights variation in carbohydrate metabolism. Together, these findings reveal extensive genetic and biochemical divergence between Indian and Southeast Asian mango groups, likely driven by historical evolutionary pressures and ecological adaptation to distinct environments.

Southeast Asian mangoes are typically polyembryonic, producing multiple embryos per seed through both sexual and asexual reproduction, a phenomenon also observed in citrus (Shukla et al., 2004; Litz, 2009). Apomictic embryos allow for clonal propagation, providing genetically uniform and vigorous rootstocks for grafting (Simon et al., 2010). A recent study reported a 3.6 kb chloroplast insertion in the promoter region of the *MiRWP* gene, which is associated with increased gene expression and the dominant polyembryonic phenotype (Yadav et al., 2023). Since this insertion is absent from current available mango reference genomes, we validated its presence by aligning PacBio HiFi sequencing reads from both polyembryonic and monoembryonic mangoes to our *M. odorata* genome assembly (Figure 6, Supplementary Table S14).

**Figure 6.**
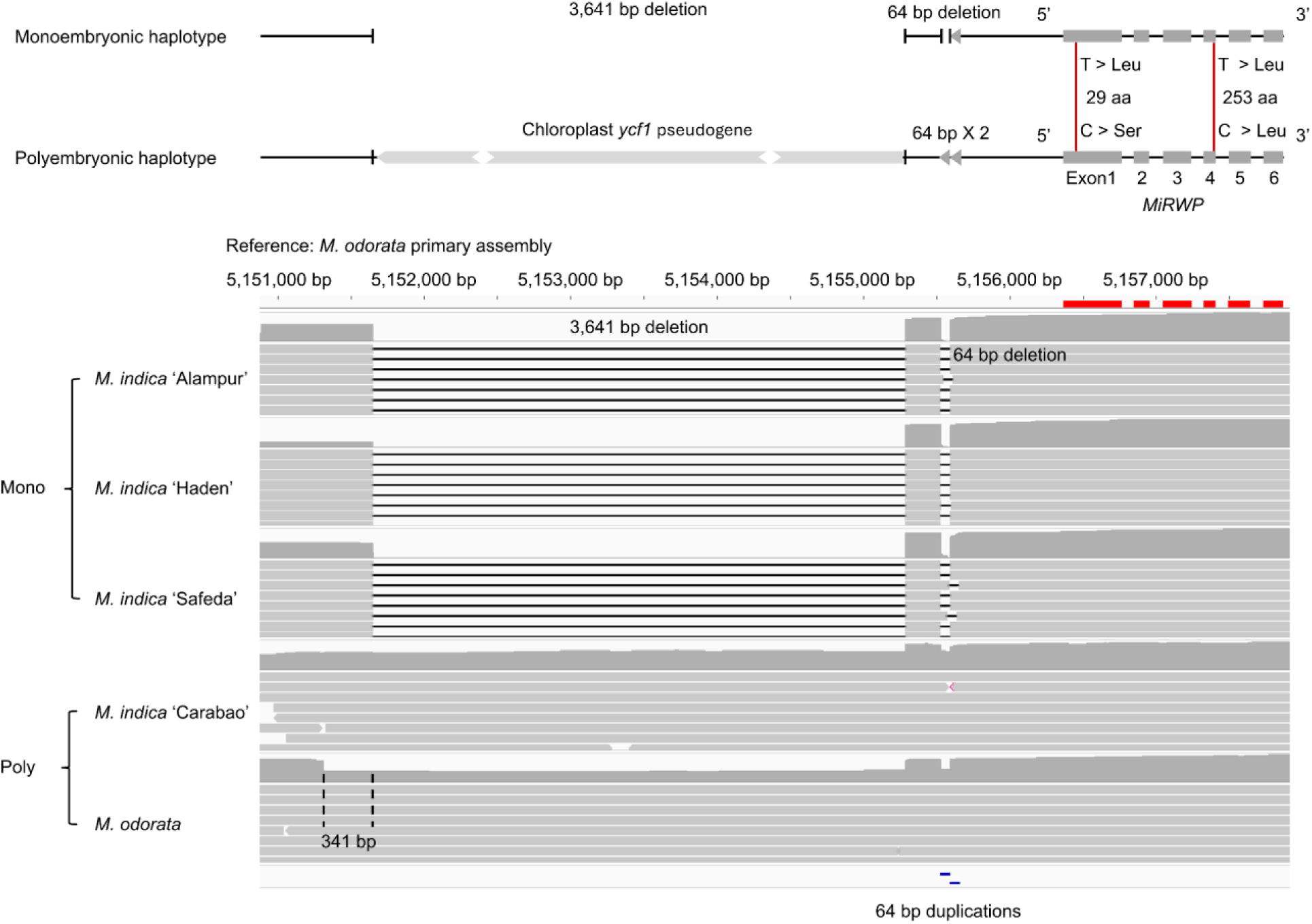
IGV visualization of HiFi read alignments from five *Mangifera* accessions to *M. odorata* chromosome 17, highlighting structural variation at the *MiRWP* promoter region. Aligned reads are represented as grey bars, missing alignments as black lines, and read depth as grey ridge plots above the alignment. The six exons of *MiRWP* are marked as red blocks. Distinct haplotype structures for monoembryony and polyembryony are illustrated. The polyembryonic haplotype contains a 3,641 bp chloroplast insertion harboring the *ycf1* pseudogene, along with two tandem 64 bp duplications upstream of *MiRWP* within the putative promoter region. In contrast, the monoembryonic haplotype lacks this 3,641 bp chloroplast insertion and carries only a single copy of the 64 bp repeat. Exon 1 and Exon 4 contain two common SNPs, indicated by red lines, with potential effects on amino acids indicated next to the lines. Monoembryonic accessions (‘Alampur’, ‘Haden’, ‘Safeda’) exhibit only the monohaplotypic genotype, while polyembryonic accessions (‘Carabao’, *M. odorata*) display a mix of polyembryonic and monohaplotypic sequences, with some variations. Short variations (<20bp) and SNPs are not shown to enhance image clarity.

The *M. odorata* genome assembly contains a 3,641 bp chloroplast-derived insertion within the putative promoter region of the *MiRWP* gene, which is associated with polyembryony in mango (Yadav et al., 2023). According to NCBI’s Conserved Domain Database, this insertion partially overlaps with the chloroplast *ycf1* gene, although substantial sequence variation suggests a possible loss of its original function. Monoembryonic accessions do not contain this insertion and also lack a 64 bp sequence downstream. In contrast, polyembryonic accessions, ‘Carabao’ and *M. odorata*, exhibit the insertion in approximately half of their sequences, accompanied by two tandem copies of the 64 bp duplication, while the remaining sequences resemble the monoembryonic haplotype. This pattern supports a dominant genetic model for polyembryony, in which the presence of a single polyembryonic haplotype—characterized by the insertions—is sufficient to confer the trait. Additionally, two common SNPs were identified within *MiRWP*: one in exon 1 results in a nonsynonymous substitution at position 29, where serine in polyembryonic haplotypes is replaced by leucine in monoembryonic ones; the second SNP, located in exon 4, is synonymous. Besides, in *M. odorata*, the sequences lacking the insertion also possess an additional 341 bp deletion upstream, a feature absent in other accessions. These findings collectively provide evidence of chloroplast DNA transfer to the nucleus, potentially altering the promoter region and influencing gene expression, thereby contributing to significant phenotypic variations related to embryony type.

## Discussion

This study investigated the genetic diversity and evolutionary relationships of mango (*Mangifera* spp.) by integrating pangenomic and population genomics analyses. The cultivated mango species *M. indica* was represented by four genetically and geographically diverse accessions, while the wild species *M. odorata* was included due to its distinct phenotypic and genetic divergence, enabling broader capture of intra- and interspecific variation. A reference-guided anchoring approach enabled high-contiguity, near telomere-to-telomere genome assemblies, providing a robust foundation for constructing a representative mango pangenome spanning two major ecotypes and multiple lineages.

Ribosomal DNA (rDNA) sequences were detected across multiple chromosomes, notably chromosomes 7, 11, and 19, as extensive tandem arrays. The repetitive and homogeneous nature of rDNA loci has long been recognized as a significant barrier in achieving complete genome assemblies in plant species (Álvarez and Wendel (2003). For example, in *Arabidopsis thaliana*, rDNA comprises hundreds of 10 kb repeat units, accounting for ∼8% of the genome (Pruitt and Meyerowitz, 1986). Although not implemented in this study, incorporating ultra-long nanopore reads exceeding 100 kb may enhance assembly contiguity in such regions (Jain et al., 2018).

The assembled mango genomes displayed similar total sizes (∼365 Mb), though the wild species *M. odorata* exhibited the highest number of predicted protein-coding genes (38,112) and a significantly higher heterozygosity rate (2.89%) compared to the cultivated mango accessions (1.41%). The results are consistent with prior reports (Li et al., 2020; Bally et al., 2021; Ma et al., 2021; Singh et al., 2021). For instance, the genome of ‘Alphonso’ was estimated at 392.9 Mb (including unplaced scaffolds), with a heterozygosity rate of 1.5% based on k-mer analysis (Wang et al., 2020), while ‘Irwin’ was assembled at 365 Mb with a heterozygosity rate of 1.24% (Wijesundara et al., 2024). The reduced heterozygosity observed in cultivated accessions likely reflects genetic bottlenecks imposed by long-term artificial selection during breeding (Hallauer and Darrah, 1985; Hyten et al., 2006).

Cultivated mangoes from Southeast Asia and India represent two ecotypes, characterized by distinct phenotypic and agronomic traits. Although these ecotypes differ markedly, molecular divergence appears relatively recent. This is illustrated by the low synonymous substitution rate (Ks = 0.0049) between Southeast Asian ‘Carabao’ and Indian ‘Safeda’, indicating either historical gene flow or parallel selection pressures across these geographically separated regions. This genetic proximity may have been influenced by trade routes, hybridization, or similar environmental adaptations during mango domestication. In contrast, Indian cultivars ‘Alampur’ and ‘Safeda’ exhibit greater divergence (Ks = 0.0067), with ‘Alampur’ exhibiting more gene family expansion than contraction, compared to the other accessions, suggesting early differentiation within the Indian gene pool (Figure 2a, 2b). This genetic disparity is likely shaped by regional geographic isolation, localized adaptation, and divergent breeding histories.

The construction of a representative pangenome, encompassing accessions from diverse genetic backgrounds, enabled a comprehensive exploration of gene family composition, shared and unique genes, and their functional implications that contributed to adaptive genetic variation. Gene family analysis showed that the total number of gene clusters plateaued after the inclusion of five to six genomes, suggesting that most gene families have been captured (Bonnici and Chicco, 2024). Core genes comprised 66.8% of total identified genes, indicating a conserved genomic framework across mango accessions. This proportion is comparable to the ∼60% reported in a pangenome study of 69 *Arabidopsis thaliana* accessions (Lian et al., 2024). Functional enrichment of core genes revealed involvement in essential biological processes such as primary metabolism, signal transduction, transcription regulation, photosynthesis, DNA repair, and intracellular transport, emphasizing their critical role in maintaining cellular and physiological functions (Kaufmann et al., 2010; Geitmann and Nebenführ, 2015; Manova and Gruszka, 2015; Stirbet et al., 2019). In contrast, dispensable and accessory genes were enriched in secondary metabolic pathways involved in biotic and abiotic stress responses, pigmentation, and ecological interactions, suggesting roles in adaptive evolution (Zaynab et al., 2018; Divekar et al., 2022; Elshafie et al., 2023; Tang et al., 2024). These findings emphasize the utility of a pangenomic framework for elucidating the genomic architecture and adaptive potential of mango.

The evolutionary history of *Mangifera* is shaped by extensive genetic diversity, wide geographic distribution, and complex domestication patterns. Our population genomic analysis revealed higher genetic diversity in Indian and Florida-USA accessions, whereas South American and Southeast Asia groups showed comparatively lower diversity and minimal admixture with other groups (Figure 4b). The observed reduced diversity in the SEA group may be influenced by limited representation in our germplasm collection, potentially introducing sampling bias. Nonetheless, the SEA accessions consistently formed a genetically distinct cluster, likely shaped by restricted germplasm exchange, localized cultivation, and domestication bottlenecks, which is consistent with previous mango genetic studies (Warschefsky and von Wettberg, 2019; Wang et al., 2020; Liang et al., 2024; Ma et al., 2024). Similarly, South American and Caribbean accessions formed a separate genetic cluster, consistent with earlier findings from Florida germplasm collections (Warschefsky and von Wettberg, 2019). However, despite this distinct clustering, the Caribbean group exhibited high genetic similarity with the Indian and Florida-USA groups, suggesting shared ancestry or historical gene flow between these regions.

Genomic differences between Southeast Asian and Indian-origin mangoes were examined through *F_st_* analysis and selection sweep comparisons. Enrichment analysis of genes near loci with high *F_st_* values revealed pathways associated with specialized metabolism, transport, and cellular processes. Notably, pathways involved in alkaloid and diterpenoid biosynthesis highlight ecological adaptations to distinct environmental pressures (O’Hagan, 2000, Matsuura and Fett-Neto, 2015; Li et al., 2021; Plazas et al., 2022). Enrichment in amino acid and lipid metabolism, including phenylalanine and alpha-linolenic acid metabolism, underscores stress response and signaling roles (Liu et al., 2021; Zi et al., 2022; Ramzan et al., 2023). Southeast Asian mangoes, originating from humid tropical rainforests, likely evolved traits for pest resistance and stress tolerance, while Indian mangoes, adapted to drier, nutrient-limited soils, may have developed mechanisms for drought resistance and nutrient uptake. However, further research is needed to confirm these ecological adaptations. These distinct ecological conditions, along with geographic and evolutionary isolation, contribute to the genetic divergence between the two major ecotypes. Mango is an important tropical crop with enriched nutrients and vitamins and holds great economic and cultural importance to farmers worldwide. However, challenges such as high heterozygosity, out-crossing, and long juvenile periods have significantly hindered its breeding process and our understanding of mango genetics. Recent advances in sequencing technology and high-throughput phenotyping offer promising solutions to these bottlenecks. The dissection of the *MiRWP* gene structural variation across different mango genomes exemplifies how pangenomic approaches can enhance our understanding of this important crop. Building on this, the present study offers valuable insights for advancing mango conservation and breeding efforts. Distinct phenotypic differences between Indian and Southeast Asian mangoes, such as size, shape, color, fiber content, and seed type reflect their divergent domestication and selection histories. Indian mangoes tend to be larger, oval-shaped, fiber-rich, and display yellow to orange hues with reddish tones upon ripening, and are typically monoembryonic. In contrast, Southeast Asian mangoes are generally smaller, more flattened, retain a green color when ripe, and are predominantly polyembryonic (Litz, 2009; Kuhn et al., 2017; Fairchild Tropical Botanic Garden, n.d.). Integrating such phenotypic information with genomic data through genome-wide association studies (GWAS), will be instrumental in identifying key functional genes underlying fruit quality traits, thereby accelerating the development of elite cultivars (Zhang et al., 2021; Zahid et al., 2022; Li et al., 2024a; Li et al., 2024b). Furthermore, expanding the mango pangenome to encompass a broader range of domesticated and wild mango relatives will not only enrich our understanding of mango evolution but also provide essential resources for targeted breeding programs.

## Methods

### Conservation and Maintenance of Mango Germplasm

The mango (*Mangifera* spp.) germplasm utilized in this study is maintained at the University of Florida Tropical Research and Education Center (UF/IFAS TREC) in Homestead, Florida, USA. The site is characterized by an equatorial monsoonal climate (Köppen classification Aw), with mean monthly temperatures ranging from 60°F to 89°F and an annual average precipitation of approximately 1,531 mm (60.3 inches) (FAWN, n.d.; WeatherSpark, n.d.). Comprehensive details regarding field design, soil properties, and management practices are available in Michael et al. (2023).

### Sample Collection and Sequencing

Four *M. indica* cultivars were selected to encompass representative genetic diversity as reported in Michael et al. (2023). These included the U.S. cultivar *M. indica* ‘Haden’; the Indian cultivars *M. indica* ‘Alampur’ (Alampur Baneshan) and *M. indica* ‘Safeda’ (Safeda Lucknow); and the Southeast Asian cultivar *M. indica* ‘Carabao’ (The Philippines). Additionally, the wild species, *M. odorata*, originating from Southeast Asia, was included to assess its phylogenetic relationship with cultivated varieties and its potential breeding value.

Young, unexpanded leaves were collected and immediately snap-frozen in liquid nitrogen. Nuclei were isolated from 1 gram of frozen tissue using the Bionano Prep Plant Tissue DNA Isolation Kit (Bionano Genomics, San Diego, CA), followed by extraction of high molecular weight (HMW) genomic DNA with the Circulomics Nanobind Plant Nuclei Big DNA Kit (Pacific Biosciences, Menlo Park, CA). DNA was sheared using a MegaRupter 3 (Diagenode, LLC, Denville, NJ) and size-selected for 20 kb fragments on a PippinHT (Sage Science, Beverly, MA). Libraries were prepared using the SMRTbell Library Prep Kit 3.0, followed by polymerase binding with the Sequel II Binding Kit 3.2, and sequenced on the Sequel IIe platform using SMRTLink v.11.0.

### Genome Assembly and Quality Evaluation

Quality control of PacBio HiFi reads was performed using FastQC v.0.12.1 (Andrews, 2010), followed by adapter trimming conducted via FCS-adaptor v.0.5.4. *De novo* Genome assemblies were generated using Hifiasm v.0.19.9 (Cheng et al., 2021) under multiple duplication purging stringencies (-l 0-3) to obtain primary contigs. Homology-based scaffolding was conducted using RagTag v.2.0.1 (Alonge et al., 2022) employing the *M. indica* ‘Irwin’ T2T reference (GWHEQCT00000000) (Wijesundara et al., 2024) serving as the guide genome.

Telomeric motifs were identified using tidk v.0.2.41 (Brown et al., 2023). Scaffold termini lacking telomeric repeats were manually extended by aligning the terminal 1,000 bp of the scaffold to a BLAST database comprising PacBio HiFi reads containing telomeric sequences (“AGGGTTT” or “AAACCCT”). To resolve internal assembly gaps, 1,000 bp sequences flanking each gap were aligned against the complete HiFi read dataset using BLAST v2.9.0 (Altschul et al., 1990). The highest-scoring alignments were selected, and the corresponding reads were used to extend the flanking sequences, thereby closing the gaps. To validate assemblies, PacBio HiFi reads were realigned to the assembly using Minimap2 v.2.28 (Li, 2018), and alignments were processed using SAMtools v.1.20 (Danecek et al., 2021). Read coverage and structural consistency were evaluated in the Integrative Genomics Viewer (IGV). GC content was assessed using QUAST v.5.2.0 (Gurevich et al., 2013) to evaluate potential contamination and biases. Heterozygosity was estimated by aligning primary and alternative haplotypes using NUCmer (MUMmer v4.0.0.beta2) (Marçais et al., 2018), extracting SNPs from non-repetitive regions with show-snps -Clr, and dividing the SNP count by the estimated genome size, following the standardized workflow (https://github.com/USDA-ARS-GBRU/StandardizedHeterozygosityEvaluation).

### Genome Annotation

*De novo* repeat libraries were constructed using RepeatModeler v.2.0 (Flynn et al., 2020). Repeat elements were subsequently identified using RepeatMasker v.4.1.1 (Smit et al., 2013–2015) by incorporating the Dfam 3.0 database (Hubley et al., 2015) alongside the custom mango repeat library derived from RepeatModeler. Alignments were performed with the rmblastn engine (provided as part of RMBlast v.2.9.0+) (Camacho et al., 2009).

Gene annotation was carried out using BRAKER v.3.0.8 (Gabriel et al., 2024), following pipeline “C”. A custom protein library was constructed by combining OrthoDB v.12.0 (Kuznetsov et al., 2022) and *M. indica* ‘Irwin’ proteins. Gene predictions were generated using GeneMark-EP+ v.4.71 (Lomsadze et al., 2005) and further refined using AUGUSTUS v.3.5.0 (Stanke et al., 2006) by incorporating the initial evidence along with additional chained CDS-part hints to enhance accuracy. Annotation statistics were computed using the agat_sq_stat_basic.pl command from AGAT v.0.4.0 (Dainat, 2020). The annotation completeness was evaluated based on the embryophyta_odb10 database using BUSCO v.5.3.0 (Manni et al., 2021).

### Gene Family and Pangenome Analysis

To investigate evolutionary relationships and functional diversification of mango, gene family analysis was performed using the primary haplotypes of five *Mangifera* accessions alongside the reference genome *M. indica* ‘Irwin’. To reduce redundancy, protein sequences were clustered at a 99% identity threshold to retain the longest isoform from each cluster using CD-HIT v.4.6.8 (Li and Godzik, 2006). Orthologous gene families were identified using OrthoFinder v.2.5.5 (Emms and Kelly, 2019).

Orthogroup distributions were visualized using Venn diagrams to illustrate shared and accession-specific gene families. The number of core and pan gene families were calculated, and a dot plot was generated to examine the relationship between the number of genomes included and the sizes of the core and pan gene sets. Stacked bar plots were used to depict gene family variability involved in the quantification and visualization of core, softcore (orthogroups present in five out of six accessions), and variable gene families.

Functional enrichment analysis was conducted to elucidate the biological roles of gene families across the six accessions. KEGG (Kyoto Encyclopedia of Genes and Genomes) pathway enrichment was performed using TBtools-II v.2.136 (Chen et al., 2023) on core and variable genes in *M. indica* ‘Irwin’ and those unique to *M. odorata.* Enriched pathways were ranked by enrichment factor and visualized as dot plots to highlight their evolutionary and adaptive significance in *Mangifera*.

### Phylogenetic Tree Construction and Whole Genome Duplication Events Analysis

Multiple sequence alignments of shared single-copy orthologs among six *Mangifera* accessions were performed using MAFFT v.7.520 (Katoh and Standley, 2013). Poorly aligned regions were removed with Gblocks v.0.91b (Talavera and Castresana, 2007) using stringent settings (-t=p, - b4=5, -b5=a). Trimmed alignments were concatenated and analyzed to infer phylogenetic relationships in IQ-TREE v.2.2.2.7.7 (Minh et al., 2020). The best-fit substitution model was selected using ModelFinder Plus (MFP) (Kalyaanamoorthy et al., 2017), and phylogenetic topology was supported by 1,000 bootstrap replicates. To minimize inflated variance, gene families exceeding 100 members were filtered out using the script clade_and_size_filter.py. Gene family expansions and contractions were analyzed using CAFE v.5.0.0 (Mendes et al., 2020) with a calibrated Newick tree and results were visualized in MEGA 11 (Tamura et al., 2021).

A cross-species phylogeny was also constructed using the concatenated single-copy orthologs. In addition to the six *Mangifera* genomes, five Sapindales species including *Pistacia vera* (GCF_008641045.1) (Zeng et al., 2019), *Pistacia integerrima* (GCA_026225825.1), *Citrus clementina* (GCF_000493195.1) (Wu et al., 2014), *Citrus sinensis* (GCF_022201045.2) (Bausher et al., 2006), and *Dipteronia sinensis* (GCA_033220585.1) (Feng et al., 2023) were included. *Vistis vinifera* (GCF_030704535.1) (Shi et al., 2023) was used as the outgroup. Divergence times were estimated using the Timetree Wizard in MEGA 11 (Mello, 2018; Tamura et al., 2021), based on divergence estimates of 78 million years between *P. vera* and *C. clementina*, and between *D. sinensis* and *P. integerrima*, as retrieved from the TimeTree 5 database (Kumar et al., 2022).

Synonymous substitution rates (Ks) for shared single-copy orthologs were calculated to assess evolutionary divergence among mango accessions. Protein alignments were performed using DIAMOND v.2.1.8 (Buchfink et al., 2014) with an e-value threshold of 1×10^−10^. Reciprocal best hits (RBH) were identified using the *dmd* function of WGD v.1.1.1 (Zwaenepoel and Van De Peer, 2019). Codon-based alignments were analyzed in PAML v.4.9h (Yang, 2007), and Ks values were computed using the *ksd* function in WGD. A Ks threshold of 0.001 was applied to filter out spurious signals. Ks distributions were visualized as ridge plots in R v.4.3.1 using ggplot2 (Wickham, 2016; R Core Team, 2023).

### Structural Variation Identification through Genome Alignment

Pairwise genome alignments between the five *Mangifera* genome assemblies and the *M. indica* ‘Irwin’ reference genome were conducted using Minimap2 v.2.28 (Li, 2018), with the alignment order guided by the inferred phylogenetic relationships. The alignment files were converted to sorted and indexed BAM format using SAMtools v.1.20. Structural variations were identified using SyRI v.1.5.4 (Goel et al., 2019). Genome-wide synteny and SVs were visualized with plotsr v.0.5.4 (Goel and Schneeberger, 2022). The SV counts were summarized and visualized as stacked bar plots using ggplot2 in R.

### Population sampling, Genotyping-by-Sequencing (GBS), and SNP calling

To investigate the genetic diversity and population structure of mango, a total of 197 accessions from *M. indica*, *M. zeylanica*, and *M. odorata* were selected from the germplasm collection located at UF/IFAS TREC (Supplementary Table S8). Additionally, one *M. casturi* accession was obtained from the Fairchild Farm collection at Fairchild Tropical Botanical Garden, Miami, FL. These accessions were sourced from over 25 countries, including the United States, India, Thailand, Vietnam, Egypt, Australia, Israel, and Trinidad.

Genomic DNA was extracted from 50 mg of fresh leaf tissue using a modified CTAB protocol (Barbier et al., 2019). Quality was assessed via 0.8% agarose gel electrophoresis and spectrophotometric absorbance ratios (260/280 nm) using a Tecan Spark® Multimode Microplate Reader. Genotyping-by-Sequencing (GBS) libraries were prepared using the restriction enzymes *NlaIII* and *MluCI* at the University of Minnesota Genomics Center and sequenced on the Illumina NovaSeq 6000 platform with an S1 flow cell (single end, 1 x 101-bp). A total of 716,368,605 reads were generated (mean of 3.64 million reads per sample), with ≥ Q30 base quality. Configuration files and trimming scripts are available at https://bitbucket.org/jgarbe/gbstrim.

Trimmed reads were aligned to the *M. indica* ‘Irwin’ reference genome (Wijesundara et al., 2024) using Burrows–Wheeler Aligner v.0.7.17 (Li and Durbin, 2009). Duplicate reads were marked with Picard v.2.9.2 (Picard Toolkit, 2019), and variant calling was performed using the HaplotypeCaller-GVCF pipeline from Genome Analysis Toolkit (GATK v.4.2.6.1) (Van der Auwera and O’Connor, 2020). Variants were filtered using thresholds: QD < 2.0, QUAL < 30.0, MQ < 20.0, MQRankSum < −12.5, and ReadPosRankSum < −8.0.

### Genomic Analysis of Population Structure

High-confidence biallelic SNPs were filtered using PLINK v.2.00 (Chang et al., 2015), retaining loci with MAF ≥ 0.05 and ≤ 10% missing data, yielding 10,482 SNPs. Population structure was inferred using ADMIXTURE v.1.23 (Alexander et al., 2009), testing hypothetical ancestral populations (*K* = 1-10). The optimal number of clusters (*K* = 4) was selected based on the lowest cross-validation error. Results for *K* = 2 to 4 were visualized as stacked bar plots using ggplot2 v.3.5.1 in R v.4.3.1. Global ancestry distributions were visualized as pie charts scaled by sample counts and overlaid on a world map using ggplot2, sf v.1.9-19 (Pebesma, 2018), rnaturalearth v.1.0.1 (Massicotte and South, 2024), and gridExtra v.2.3 (Auguie and Antonov, 2017).

Principal component analysis (PCA) was conducted with PLINK v.2.00, and the first two principal components were plotted to illustrate genetic structure, colored by ADMIXTURE clusters (*K* = 4). SNP data were converted from VCF to PHYLIP alignment format using vcf2phylip.py v.2.8 (Ortiz, 2019), and a maximum likelihood phylogenetic tree was constructed using IQ-TREE v.2.2.2.7 with 1,000 bootstrap replicates and automatic model selection (-m MFP). The resulting tree was visualized using the Interactive Tree of Life (iTOL) v.6 (Letunic and Bork, 2024), with branches color-coded to represent the four ancestry clusters, aiding the interpretation of evolutionary relationships.

Linkage disequilibrium (LD) decay within each ancestry cluster was evaluated by calculating *r*^2^ values for 10,482 SNP pairs using PopLDdecay v.3.40 (Zhang et al., 2018). The results were grouped into distance bins with step sizes of 1,000 bp and 20,000 bp (with a breakpoint at 10,000 bp), extending up to 3 Mb, and visualized as LD decay curves using Plot_MultiPop.pl and ggplot2 in R. Genetic differentiation among the four clusters was quantified using the fixation index (*F*_ST_), calculated with VCFtools v.0.1.16 (Danecek et al., 2011).

### Analysis of Genomic Differentiation and Selection

Given the close genetic relationship among the U.S., Caribbean, and Indian populations and their likely shared origin in India, these groups were merged into a single Indian-origin (IND) group for comparative analysis against the Southeast Asian (SEA) group. Genetic differentiation and diversity were assessed by calculating pairwise *F*_ST_ values using VCFtools v.0.1.16, and nucleotide diversity (π) was estimated within each group independently using a 200 kb window and 50 kb step to capture regional variation. Genomic regions under potential selection were identified using SweeD v.3.3.2 (Pavlidis et al., 2013), with 10 kb windows for high-resolution detection. Results were visualized as Manhattan and sliding window plots using the qqman package v.0.1.9 (Turner, 2018) in R, with grey dashed lines marking the 99.9th percentile threshold to highlight candidate selective sweeps.

KEGG pathway enrichment analysis was performed on genes located within 200 kb windows (±100 kb from SNPs) centered on the top 5% of *F*_ST_ loci, to identify functional pathways associated with genetic differentiation between groups. The 200 kb window size was chosen based on an estimated linkage disequilibrium decay threshold (*r^2^*≈0.2), allowing capture of relevant candidate genes while minimizing unrelated genomic regions. This approach has been applied in previous studies (Dujak et al., 2023; Li et al., 2024a; Li et al., 2024b) to investigate genes underlying fruit quality traits.

## Author Contributions

J.L. conducted the genome assembly, performed all formal data analysis and visualization, and wrote the manuscript. V.N.M. performed sample preparation, constructed the GBS library, and generated data. S.C. assisted with the genome assembly. S.A.S., R.C.Y., B.E.S., and A.M.H.-K. handled DNA extraction, sequencing library preparation, and data generation. J.C. maintained the mango germplasm and reviewed the manuscript. A.M.H.-K. edited the manuscript, contributed to the conceptual design, and provided funding. X.W. conceptualized the original design, supervised the project, edited the manuscript, facilitated collaboration, and provided funding.

## Acknowledgments

This work was supported by the United States Department of Agriculture - Agricultural Research Service CRIS Project Number 6066-21310-006-00D and grant number 58-6066-9-006.

## Data Availability

The PacBio HiFi sequencing data and whole-genome assemblies have been deposited in NCBI under BioProject ID PRJNA1214767, with corresponding Biosample accessions SAMN46384120, SAMN46384121, SAMN46384122, SAMN46384123, and SAMN46384124. All code used in this study is available on GitHub at https://github.com/UF-Wu-Lab/Jin.

## Supporting information

Supporting information is available online in the supplementary material documents.

**Figure S1** Cross-validation error for population admixture analysis of 197 mango accessions at UF/IFAS TREC.

**Figure S2** Linkage disequilibrium (LD) decay in four mango subpopulations across 197 accessions at the UF/IFAS TREC field.

**Figure S3** KEGG enrichment of genes associated with high *F_ST_* SNPs between Indian and Southeast Asian groups.

**Table S1** PacBio HiFi sequencing statistics.

**Table S2** Repeat annotation and gene features of five *Mangifera* assemblies.

**Table S3** Orthogroup distribution across core, softcore, shell, and cloud components in *Mangifera* accessions.

**Table S4** KEGG enrichment analysis of Irwin genes in core gene clusters shared by five *Mangifera* indica cultivars and the wild-type *Mangifera* odorata.

**Table S5** KEGG enrichment analysis of Irwin genes in variable gene clusters among five *Mangifera* indica cultivars and the wild-type *Mangifera* odorata.

**Table S6** SyRI (Synteny and Rearrangement Identifier) results for five mango assemblies aligned to the reference genome of accession Irwin.

**Table S7** KEGG enrichment analysis of unique genes in *Mangifera* odorata compared to five *Mangifera* indica cultivars (Alampur, Safeda, Irwin, Haden, Carabao).

**Table S8** Population analysis of 197 mango accessions at the University of Florida Tropical Research and Education Center (UF TREC).

**Table S9** Dataset of 10,482 SNP markers from 197 mango accessions at UF TREC.

**Table S10** FST analysis between the combined Indian Group and Southeast Asian group showing differentiation based on 10,482 SNP markers from 197 mango accessions at UF TREC.

**Table S11** Nucleotide diversity (π) across 10,482 SNPs in the combined Indian group and Southeast Asian group, calculated using a 200 kb window and 50 kb step size.

**Table S12** SweeD analysis of genome-wide selective sweeps in the combined Indian and Southeast Asian groups, based on 10,482 SNPs from 197 mango accessions at UF TREC (10 kb window Size).

**Table S13** KEGG enrichment of genes identified within 200 kb windows around the top 5% *F_ST_* loci, highlighting divergence between Indian and Southeast Asian groups.

**Table S14** Genomic Coordinates and Sequences of *MiRWP* in M. odorata Reference Genome.

